# GRHL2 enhances phosphorylated estrogen receptor DNA-binding and regulates ER-mediated transcriptional activation and repression

**DOI:** 10.1101/2022.05.19.492733

**Authors:** Rebecca M. Reese, Kyle T. Helzer, Kaelyn O. Allen, Christy Zheng, Natalia Solodin, Elaine T. Alarid

## Abstract

Phosphorylation of estrogen receptor α (ER) at serine 118 (pS118-ER) is induced by estrogen and is the most abundant post-translational mark associated with a transcriptionally active receptor. Cistromic analysis of pS118-ER from our group found enrichment of the GRHL2 motif near pS118-ER binding sites. In this report we use cistromic and transcriptomic analyses to interrogate the relationship between GRHL2 and pS118-ER. We found that GRHL2 is bound to chromatin at pS118-ER/GRHL2 co-occupancy sites prior to ligand treatment, and GRHL2 binding is required for maximal pS118-ER recruitment. pS118-ER/GRHL2 co-occupancy sites were enriched at active enhancers marked by H3K27ac and H3K4me1, along with FOXA1 and p300. Transcriptomic analysis yielded four subsets of ER/GRHL2 co-regulated genes revealing that GRHL2 can both enhance and antagonize E2-mediated ER transcriptional activity. Gene ontology analysis identified several coregulated genes involved in cell migration. Accordingly, knockdown of GRHL2 combined with estrogen treatment resulted in increased cell migration but no change in proliferation. These results support a model in which GRHL2 binds to select enhancers and facilitates pS118-ER recruitment to chromatin which then results in differential activation and repression of genes that control ER-positive breast cancer cell migration.

## INTRODUCTION

Nuclear receptors are a class of transcription factors that modulate gene expression through DNA interactions in response to stimuli such as steroid hormones. One such receptor is estrogen receptor α (ER), which is expressed in over 70% of all breast cancers and is a major therapeutic target (1–3). ER transcriptional activity is largely governed by the binding of its ligand, estrogen (E2), to the C-terminus of the receptor (4, 5). Subsequently, the receptor binds to its consensus sequence, the estrogen response element (ERE; 5′-GGTCAnnnTGACC-3′) (6, 7), and interacts with other coregulators to modulate specific gene expression (8–10).

ER transcriptional activity can be altered by a variety of post-translational modifications (PTMs), including phosphorylation of several residues (11). Of particular note is phosphorylation of the serine 118 residue (pS118-ER) in the N-terminus of the receptor which can be induced in both a ligand-dependent and independent manner (12, 13). Clinically, pS118-ER is associated with low grade, well-differentiated breast tumors (14, 15). Mutational analysis assessing the function of pS118 has found the presence of pS118 is required for maximal E2-mediated ER transcriptional activity (16–18). Our group recently completed a genome-wide analysis of pS118-ER DNA-binding sites and found pS118-ER preferentially binds enhancers marked by H3K27ac in breast cancer cells (19). In addition, motif analysis comparing sites bound by pS118-ER and ER without pS118 identified enrichment of the GRHL2 motif (5’-AACCGGTT-3’) (20–23) near pS118-ER binding sites (19). GRHL2 is a member of the Grainyhead-like protein family (GRHL), a class of transcription factors which regulate epithelial differentiation (24–34). GRHL2 has been implicated in a number of cancers, demonstrating both tumor suppressive and oncogenic roles (35–54). The genomic analysis of pS118-ER DNA binding provided early evidence of interplay between GRHL2 and ER in breast cancer cells.

There is increasing evidence that GRHL2 plays a role in facilitating ER transcriptional activity. GRHL2 was identified as an ER interactor in rapid immunoprecipitation mass spectrometry of endogenous protein (RIME) studies (55), and this interaction increases upon estrogen treatment (56). Cistromic studies of ER and GRHL2 found that 29-35% of ER binding sites are co-occupied by a GRHL2 binding site (57, 58), indicating these two factors may function together to regulate gene transcription. Studies suggest that GRHL2 acts through enhancer regions to facilitate the recruitment of MLL3, a lysine methyltransferase responsible for depositing the enhancer marker H3K4me1 (58, 59). GRHL2 also co-occupies chromatin with FOXA1, a pioneer factor associated with ER transcription, which promotes an open chromatin structure and is found in regions marked by H3K4me1 (58, 60–63). GRHL2 chromatin-binding can also be enhanced by E2 (19, 56) and E2-responsive GRHL2 binding sites are found more frequently in enhancer regions identified by ER ChIA-Pet (56). Together with our work identifying the enrichment of the GRHL2 motif near pS118-ER binding sites, this evidence supports a role for co-regulation of gene expression by GRHL2 and ER at enhancer regions. We also found pS118-ER preferentially binds near genes that are up-regulated by E2, suggesting GRHL2 may co-activate genes with pS118-ER. However, GRHL2 can also suppress the production of enhancer RNAs (eRNA) at ER enhancers for *GREB1*, *TFF1*, and *XBP1* (56), indicating that GRHL2 is also capable of repressing ER transcriptional activity. The exact nature of the role of GRHL2 in modulating ER transcriptional activity, more specifically pS118-ER transcriptional activity, and the biological processes co-regulated by both factors are currently unclear.

In this report we examine the coordinate activities of GRHL2 and pS118-ER in the regulation of gene expression in breast cancer cells using cistromic and transcriptomic approaches. Our data show GRHL2 has a regulatory role in pS118-ER DNA binding at active enhancers, and GRHL2 can both activate and repress pS118-ER target genes. We also found the combined activities of GRHL2 and pS118-ER control the migratory behavior of breast cancer cells. These studies indicate a specific role for GRHL2 in modulating pS118-ER transcriptional activity.

## MATERIALS AND METHODS

### Cell culture

MCF7, BT-474, SK-BR-3, MDA-MB-468, and MDA-MB-231 cells were maintained in 10% CO_2_ at 37ᵒC with high-glucose (4.5 g/L) Dulbecco’s modified Eagles Medium (DMEM; Thermo Fisher, #11965092) supplemented with 10% HyClone fetal bovine serum (FBS; Cytiva, #SH30910.03), 100 U/mL penicillin 100 ug/mL streptomycin (1% Pen/Strep; Gibco, #15140-122), and 1 mM sodium pyruvate (Gibco, #11360-070). T47D and ZR-75-1 cells were maintained in Roswell Park Memorial Institute 1640 (RPMI; Thermo Fisher, #11875-093) supplemented with 10% FBS and 1% Pen/Strep. BT-20 cells were maintained in Minimum Essential Medium Alpha (MEMα; Thermo Fisher, #41061-029) supplemented with 10% FBS and 1% Pen/Strep.

### Protein Preparation and Western Blot

Protein samples were lysed in 1X Sample Buffer (62.5 mM Tris pH6.8, 2% SDS, 10% glycerol, 5% β-mercaptoethanol) and subsequently boiled at 100ᵒC for 10 minutes. Protein concentration for each sample was quantified using the RC DC protein assay (Bio-Rad, #5000122) and 30-50 μg of protein were run on a 10% SDS-PAGE gel. Proteins were transferred to a PVDF membrane (VWR, Cytiva, #10061-492), blocked with 5% powdered milk in TBST (50 mM Tris-HCl pH7.4, 150 mM NaCl, 0.1% Tween-20), and incubated overnight at 4ᵒC in primary antibody. Primary antibodies used were ER (Santa Cruz Biotechnology, #HC-20), pS118-ER (Abcam, #ab32396), GRHL1 (Santa Cruz Biotechnology, #sc-515541), GRHL2 (Millipore Sigma, #HPA004820), GRHL3 (Santa Cruz Biotechnology, #sc-398838), β-actin (Millipore Sigma, #A5441), and TBP (Thermo Fisher, #MA1-21516). Blots were incubated in secondary antibody for one hour at room temperature (anti-rabbit; Amersham, #NA934) (anti-mouse; Amersham, #NA931) and visualized with Clarity Western ECL Substrate (Bio-Rad, #1705061) on a ChemiDoc Imaging System (Bio-Rad).

### Drug Treatments

Prior to all drug treatments, cells were deprived of estrogens by culturing them in phenol-red free DMEM or RPMI supplemented with 10% charcoal stripped FBS (ssDMEM/ssRPMI; Cytiva, #SH30910.03, stripped in-house), 1% Pen/Strep, 2 mM L-glutamine (Thermo Fisher, #25030-081), and 1 mM sodium pyruvate (DMEM only) for three days. Cells were treated with vehicle (0.1% EtOH) or 10 nM 17β-estradiol (E2; Millipore Sigma #E2257). The duration of each treatment is noted for each experiment.

### Knockdown of GRHL2 with siRNA

For experiments involving GRHL2 knockdown, cells were reverse transfected with 25 nM of either All Stars Negative control siRNA (siCtrl; QIAGEN, #1027280), siGRHL2 #1 (RNA-seq and qPCR validation experiments) (Santa Cruz Biotechnology, #sc-77606), or siGRHL2 #2 (qPCR validation) (Dharmacon, #L-014515-02-0050) using 2.5 ug/mL of Lipofectamine 2000 (Thermo Fisher Scientific, #11668019) in Pen/Strep-free media.

### RNA Isolation and RT-qPCR

RNA isolation was performed using the QIAGEN RNeasy Mini Kit (#74106) including treatment with DNase (QIAGEN, #79254). Prior to qPCR, cDNA was generated using 1 μg of RNA and the iScript cDNA Synthesis Kit (Bio-Rad, #1708891). qPCR was performed using iQ SYBR Green Supermix (Bio-Rad, #1708884). Absolute quantification of RNA levels was performed by creating standard curves with plasmids for *GRHL1* (Origene, #RC229312), *GRHL2* (Origene, #RG214498), and *GRHL3* (Origene, #RC213012). The primers used were: *GRHL1*; Forward: 5’-ACAGCAAAAGAAACAGCATACCA-3’ Reverse: 5’-TTGCCATGGGGAAGGACAAT-3’. *GRHL2*; Forward: 5’-GCCACCAAATCTCTCCGTCA-3’ Reverse: 5’-CCACCATCACCACACTCCTG-3’. *GRHL3*; Forward: 5’-TGCAAGCGAGGAATCTTAGTCA-3’ Reverse: 5’-CTCGAGAGGCCTTACAGCTC-3’. Relative RNA levels were calculated using the ΔΔCt method. The primers used were: *LYPD3*; Forward: 5’-TCACCTTGACGGCAGCTAAT-3’ Reverse: 5’-GAACCCTGGGCCTGTTACTC-3’. *EFEMP1*: Forward: 5’-TTATCATGGCGGCTTCCGTT-3’ Reverse 5’-TGGGCAAACACATCGGTTCT-3’. *ARHGEF28*: Forward: 5’-TGGAAAACTGCTACAGGTCGT-3’ Reverse 5’-ATGGCTTCTGATCAACGGCT-3’. *SCUBE2*: Forward: 5’-TTCACCGCTCGGAAGAGGG-3’ Reverse: 5’-TGGTTCTTGGCCAGCTCAAA-3’. *RPLP0* (housekeeping gene): Forward: 5’-GACAATGGCAGCATCTACAAC-3’ Reverse: 5’-GCAGACAGACACTGGCAAC-3’.

### RNA-seq

MCF7 cells and T47D cells were cultured in 10 cm plates in ssDMEM and ssRPMI, respectively, for two days. On day three, cells were transfected as described above at a density of 200,000 cell/well in 12-well plates in Pen/Strep-free estrogen-deprived media. After 24 hours, media was changed to estrogen-deprived media containing Pen/Strep and cells were treated with E2 or vehicle for 24 hours. RNA was then isolated as described above. Samples for qPCR validation of RNA-seq were prepared using the same process. Sequencing libraries were prepared using the Illumina TruSeq Stranded mRNA Library Prep kit (Illumina, #20020595) and sequenced for 150bp paired-end reads on the NovaSeq6000 by the Next Generation Sequencing department at the UW-Madison Biotechnology Center.

Quality control assessment of fastq files was performed using FASTQC (64). Adaptor trimming to remove Illumina adaptors was performed using Cutadapt (65) through Trim Galore (66). Reads were aligned to the hg38 genome with STAR (67). Aligned reads were assigned to genes using featureCounts from the Subread package (68) and the hg38 NCBI RefSeq GTF (available at (69)) Differential gene expression was performed using the DESeq2 package (70) with fold change shrinkage using ashr (71). PCA plot was generated using DESeq2, heatmaps generated using pheatmap (72), and gene ontology was performed using clusterProfiler (73).

### Migration Assays

Migration assays were performed by seeding estrogen-deprived wild-type and CRISPR GRHL2 knockout cell pools obtained from Synthego (Menlo Park, CA) at a density of 700,000 cells/mL in Ibidi 4-well dishes (Ibidi, #80406) simultaneously with vehicle or E2 treatment. 24 hours later, the 4 well insert was removed to create a cell-free gap. The plates were imaged at 0 hours, 6 hours, and 24 hours, and the gap was measured using the Wound Healing Size Tool plugin for imageJ (74).

### Cell Cycle Analysis

Cells were deprived of estrogens for three days prior to adding treatments as described above for 24h, 48h, and 72h. Cell pellets were fixed in 100% ethanol and then resuspended in a staining solution consisting of 100 ug/mL RNase A (Invitrogen, #12091021) and 50 ug/mL propidium iodide (Millipore Sigma, #P4170). The dye intensity of 10,000 events was captured on a Thermo Fisher Attune NxT Flow Cytometer. Analysis was performed using the ModFit software.

### CUT&RUN

MCF7 cells were cultured in 10cm plates in ssDMEM for one day. On day two, cells were transfected as described above at a density of 350,000 cell/well in 6-well plates. After 24 hours, the media was changed to the appropriate media containing Pen/Strep. The following day, cells were removed from the plates using brief trypsinization and assessed for viability using trypan blue (Gibco, #15250-061). 5×10^5^ viable cells were treated with vehicle or E2 for 30 min while rotating at 37ᵒC. Following treatment, samples were processed for CUT&RUN using the CUTANA ChIC/CUT&RUN Kit (Epicypher, #14-1048; protocol from user manual version 2.0). Antibodies used were ER (Epicypher, #13-2012), pS118-ER (Santa Cruz Biotechnology, #sc-12915), GRHL2 (Millipore Sigma, #HPA004820), and IgG (Negative control; Epicypher, #13-0042).

DNA concentrations were measured using the Qubit dsDNA HS assay kit (Thermo Fisher, #Q33231). Library prep was performed using the NEBNext Ultra II DNA Library Prep Kit for Illumina (NEB, #E7103) and NEBNext Multiplex Oligos for Illumina (NEB, #E6440), with changes made to the protocol to improve the capture of smaller TF fragments: the inactivation temperature after end-repair was changed to 50°C for 1 hour, the bead size-selection ratio in the adaptor cleanup step was increased to 1.75x, and the bead size-selection ratio in the PCR cleanup step was increased to 1.2x (75, 76). Samples were sequenced by paired-end 150bp sequencing on an Illumina NovaSeq 6000 by the Next Generation Sequencing department at the UW-Madison Biotechnology Center.

Quality control assessment and adaptor trimming was performed as described in “RNA-seq”. Reads were aligned to the hg38 genome using Bowtie2 (77) using the --very-sensitive-local and --dovetail flags. Aligned fragments were filtered for a maximum fragment length of 120bp using alignmentSieve from Deeptools (78). Bam files from replicates were merged using Samtools merge. Peaks were called with MACS2 (79) using IgG as a control and an FDR cutoff of 0.05. Peak overlaps were performed with HOMER mergePeaks (80) with -d given. Heatmaps were plotted using Deeptools computeMatrix and plotHeatmap.

### Generation of S118A ER mutation in MCF7 Cell Line

Insertion of the S118A ER mutation in MCF7 cells was performed by following the Alt-R™ CRISPR-Cas9 System Protocol (IDT; Coralville, IA, USA) for use with the Neon™ Transfection System (Thermo Fisher, #MPK5000) with alterations specific for MCF7 cells. MCF7 cells were brought up at a low passage (p2) as recommended by the electroporation protocol. Cells were grown in DMEM supplemented with 10% FBS and 1% Pen/Strep, split once, and then grown to confluency prior to cell counting and electroporation. The crRNA:tracrRNA duplex was formed by resuspending the Alt-R™ CRISPR-Cas9 crRNA (5’ – TGCAGGAAAGGCGACAGCTG – 3’) and Alt-R™ CRISPR-Cas9 tracrRNA (IDT, #1072532) in equimolar concentrations in nuclease-free IDTE buffer (10 mM Tris, 0.1 mM EDTA, pH 7.5; IDT, #11-01-02-02) to a final duplex concentration of 44 µM. The mixture was brought to 95°C for 5 min and allowed to cool to room temperature. To form the RNP complex, Alt-R™ S.p. Cas9 Nuclease 3NLS (IDT, #1074181) was resuspended in Resuspension Buffer R (Thermo Fisher) to a final concentration of 36 µM. For each electroporation, equal volumes of the crRNA:tracrRNA duplex and RNP complex were combined and incubated at room temp for 20 min. Prior to electroporation, MCF7 cells were collected, washed with 1xPBS, counted, and resuspended in Resuspension Buffer R to a concentration of 5 x 10^6^ cells/mL. 8 µL of the cell suspension was combined with the crRNA:tracrRNA:RNP complex, ssODN donor (5 –CCCCCACTCAACAGCGTGTCTCCGAGCCCGCTGATGCTACTGCACCCGCCGCCGCAATTGGCGCCGTTCCTGC-AGCCCCACGGCCAGCAGGTGCCCTACTACCTGGAGAACGAGCCCAGC – 3’), and Alt-R™ electroporation enhancer (IDT, #1075915) for final concentrations of 1.8 µM crRNA:tracrRNA, 1.5 µM Cas9 RNP complex, 4 µM ssODN donor, and 1.8 µM electroporation enhancer. As a control, MCF7 cells with no crRNA:tracrRNA were separately transfected keeping the same concentrations of the other components. The final cell suspensions were electroporated using the Neon™ Transfection System with parameters set for MCF7 cells (2 pulses, pulse voltage 1250v, pulse width 20ms). Cells were transferred to 12-well plates with pre-warmed media without Pen/Strep and allowed to recover for 72 h.

### Screening of CRISPR-Cas9 clones

After recovery, the cells were grown to confluency and cell populations were screened for presence of the S118A mutation. DNA was isolated using the QIAGEN DNeasy Blood & Tissue Kit (QIAGEN, #69504) and primers were generated to amplify around the region of interest (Forward: 5’ – GCCAACGCGCAGGTCTAC – 3’; Reverse: 5’ – TCAAAGCGCCCCGTGTTTAT – 3’). Amplicons were subject to digestion by SfoI (NEB, #R0606S) to test for the insertion of the S118A mutation. Populations positive for SfoI digestion, indicating insertion of S118A mutation, underwent limiting dilution to identify single colonies. Colonies were selected and expanded, followed by PCR screening and SfoI digestion as described above. PCR amplicons from positive clones were additionally sequenced to confirm insertion of the S118A mutation. Positive clones were further validated by induction of pS118-ER by E2 by Western blot analysis, as described above.

### Chromatin Immunoprecipitation Sequencing (ChIP-seq)

Cells were plated at a density of six million cells per 10cm plate with two plates per condition and were deprived of estrogen for three days, followed by a 30-minute treatment with vehicle or E2. Samples were prepared for ChIP as previously described (19) using a GRHL2 antibody (Millipore Sigma, #HPA004820). DNA concentrations were measured using the Qubit dsDNA HS assay kit (Thermo Fisher, #Q33231). Library prep was performed using the NEBNext Ultra II DNA Library Prep Kit for Illumina (NEB, #E7103) and NEBNext Multiplex Oligos for Illumina (NEB, #E6440). Samples were sequenced by paired-end 150bp sequencing on an Illumina NovaSeq 6000 by the Next Generation Sequencing department at the UW-Madison Biotechnology Center.

Quality control assessment and adaptor trimming was performed as described in “RNA-seq”. Reads were aligned to the hg38 genome with Bowtie2 using --local and peaks were called with MACS2 using an FDR of 0.01. All peak file merging was performed using HOMER mergePeaks, -d given. For ChIP-seq data from previous publications, data was downloaded using the SRA-Toolkit (81) from the following accession numbers: pS118-ER/ER ChIP-seq (GSE117569) (19), GRHL2 ChIP-seq (GSE109820) (56), FOXA1 ChIP-seq (GSE59530) (82), p300 ChIP-seq (GSE60270) (83), H3K4me1/3 and H3K27ac ChIP-seq (GSE57436) (84), and H3K27me3 ChIP-seq (GSE108787) (85). All data was trimmed and aligned using the previously described methods with appropriate changes for single-end data. pS118-ER/ER and GRHL2 peaks were called using the narrowpeak setting in MACS2 with FDR cutoffs of 0.01 and 0.05, respectively. Motif analysis was performed using annotatePeaks.pl and findMotifsGenome.pl from HOMER.

### Statistical Methods

The Wilcoxon rank sum test was used to analyze differences in RNA levels and percent gap closure. A p-value less than 0.05 was considered significant.

### Data Availability

The data reported in this paper have been deposited in the Gene Expression Omnibus (GEO) database, www.ncbi.nlm.nih.gov/geo, under accession number GSE202622.

## RESULTS

### GRHL2 expression is increased in ER-positive breast cancer compared to ER-negative

The GRHL family is composed of three members, GRHL1, GRHL2, and GRHL3, all of which regulate gene expression via the same DNA-binding motif (5’-AACCGGTT-3’). To characterize the GRHL family in breast cancer, we first assessed the RNA expression of the GRHL family in normal breast tissue and breast tumors using GEPIA2 (86). *GRHL2* expression was significantly increased in tumors than in normal breast, whereas *GRHL1* and *GRHL3* showed no significant differences (Fig. 1A). These observations were further supported by examination of TCGA datasets via cBioportal which showed that *GRHL2* was amplified in 10.8% of breast cancers while *GRHL1* and *GRHL3* were amplified in less than 1% of breast cancers (54, 87–90).

**Figure 1:**
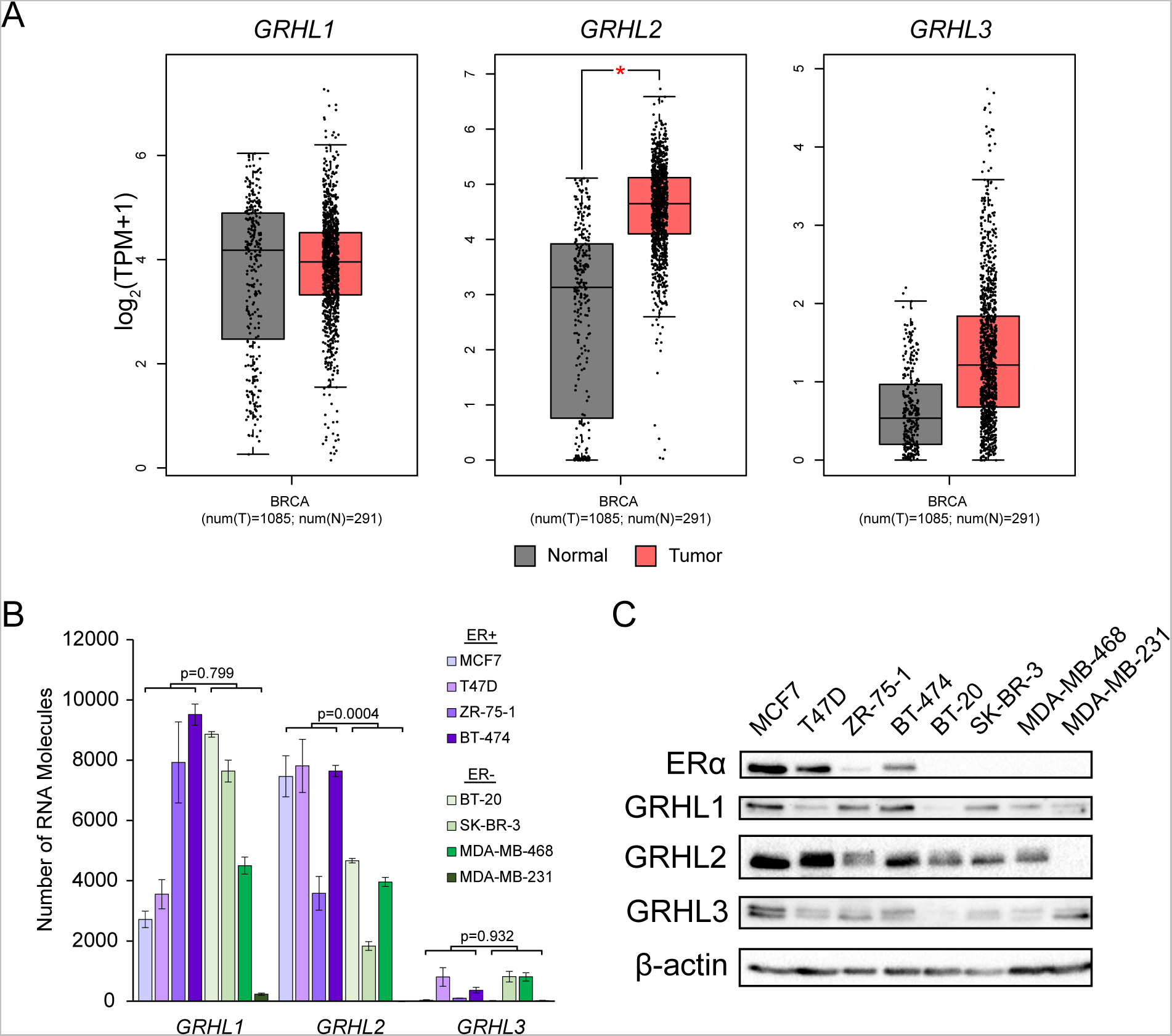
GRHL2 expression is higher in ER-positive vs ER-negative breast cancer. A) mRNA expression of the *GRHL* family in normal and tumor breast tissue obtained from GEPIA2. A * indicates a p-value < 0.05. B) Absolute quantification of *GRHL* mRNA and in a panel of ER-positive and ER-negative breast cancer cell lines. Error bars represent mean ± standard error. C) Representative Western blot of GRHL protein levels in the same cell panel. β-actin is used as a loading control.

An important consideration in treating breast cancer is the ER status of the tumor. We assessed protein and RNA expression of the GRHL family in four ER-positive breast cancer cell lines (MCF7, T47D, ZR-75-1, BT-474), and four ER-negative cell lines (BT-20, SK-BR-3, MDA-MB-468, MDA-MB-231). Each of the GRHL family members is expressed in all evaluated cell lines, with the exception of MDA-MB-231 which lacks GRHL2 expression (Fig. 1B,C). *GRHL2* mRNA expression was significantly higher in ER-positive than ER-negative cell lines, whereas *GRHL1* and *GRHL3* expression did not differ significantly between the two categories (Fig. 1B). These results were further corroborated by a representative immunoblot of the GRHL family in breast cancer cell lines (Fig. 1C). A previous study in primary breast tumors also found a significant correlation between GRHL2 and ER expression (40). Given the proclivity for GRHL2 expression in ER-positive breast cancer, we focused solely on GRHL2 for the remainder of our studies.

### pS118-ER/GRHL2 co-occupancy sites are found in enhancer regions

In previous work from our group, we noted an enrichment of the GRHL2 motif near sites occupied by ER phosphorylated at serine 118 (pS118-ER) (19). We asked whether there were differences in the binding signals of pS118-ER/GRHL2 co-occupancy sites compared to those where each factor occupies chromatin independently of the other. We took advantage of our own previously published pS118-ER and ER ChIP-seq datasets (19) and a GRHL2 ChIP-seq performed in MCF7 cells treated with E2 for 30 and 45 minutes, respectively (56). The pS118-ER binding sites consist of a subset of sites within the total ER binding sites. ER binding sites are defined as sites within the total ER binding set, excluding those that are bound by pS118-ER. We identified 4,192 pS118-ER/GRHL2 co-occupancy sites and 4,028 ER/GRHL2 co-occupancy sites which comprise 28% and 12% of total pS118-ER and total ER binding sites, respectively. Of the binding sites occupied independently by each factor, we identified 33,797 ER sites (ER-only), 10,842 pS118-ER sites (pS118-ER-only), and 12,150 GRHL2 sites (GRHL2-only) (Fig. 2A). Of the co-occupancy categories, GRHL2 binding was found prior to ligand treatment at 82.8% of the pS118-ER/GRHL2 co-occupancy sites and 83.2% of the ER/GRHL2 co-occupancy sites, indicating GRHL2 can bind the majority of these regions before E2-induced recruitment of ER (Fig. 2B). An example of a pS118-ER/GRHL2 co-occupancy site is displayed in Fig. 2C. These results confirm GRHL2 is enriched at pS118-ER binding sites relative to ER and indicate that GRHL2 and ER have both dependent and independent functions.

**Figure 2:**
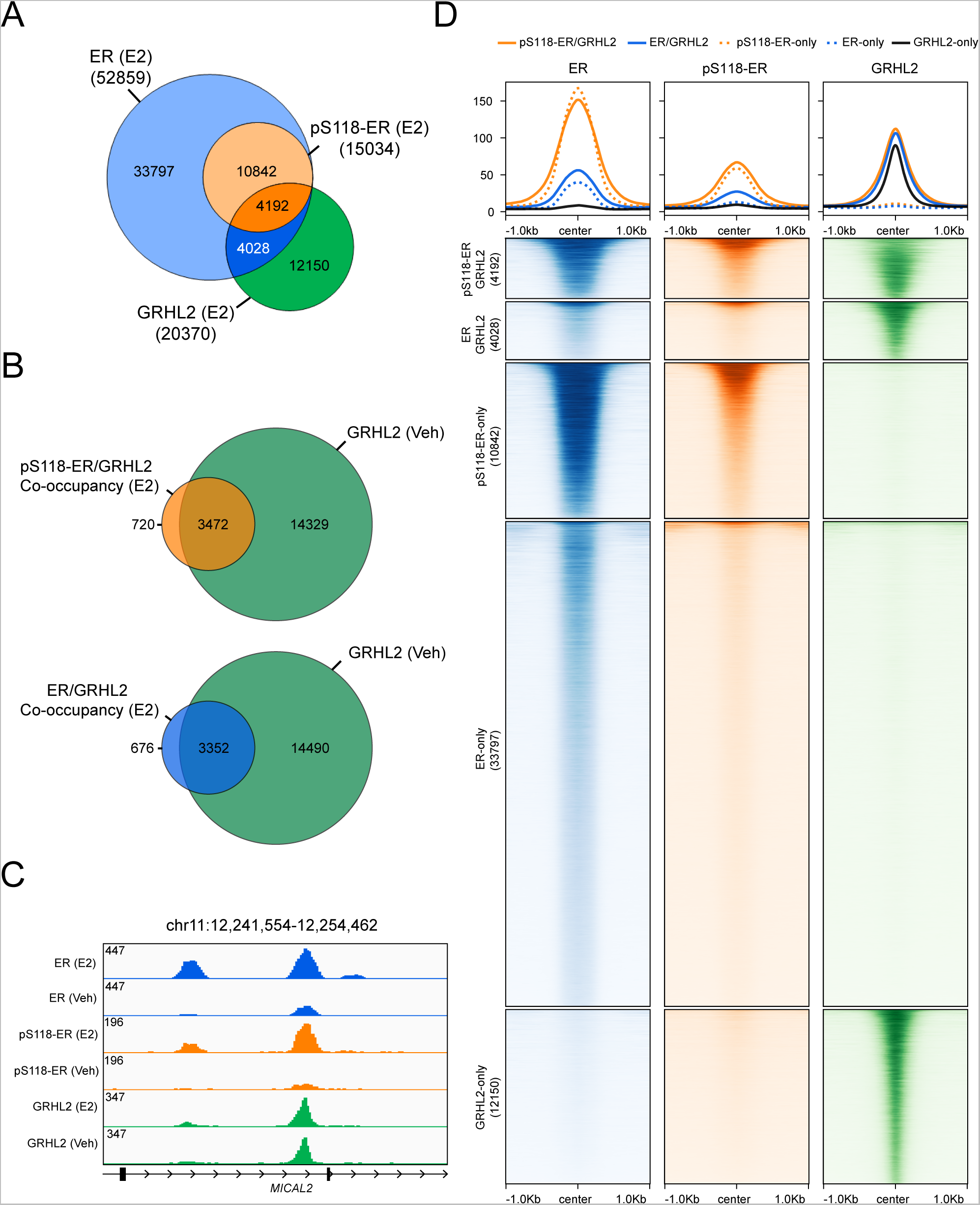
Identification of pS118-ER/GRHL2 co-occupancy sites. A) Venn diagram showing the overlap of MCF7 E2-treated total ER, pS118-ER, and GRHL2 ChIP-seq datasets. pS118-ER binding sites are a subset of total ER bindings sites. ER sites are defined as those that do not overlap with the pS118-ER dataset. Five binding categories were established: pS118-ER/GRHL2 co-occupancy (4,192 sites), ER/GRHL2 co-occupancy (4,028 sites), pS118-ER-only (10,842 sites), ER-only (33,797 sites), and GRHL2-only (12,150 sites). B) Venn diagrams showing the overlap of pS118-ER/GRHL2 co-occupancy sites and ER/GRHL2 co-occupancy sites with GRHL2 peaks detected in vehicle conditions. C) Representative genome browser track of a pS118-ER/GRHL2 co-occupancy site. D) Heatmaps showing binding signal of ER, pS118-ER, and GRHL2 at each of the five binding categories. Data were obtained from previously published ChIP-seq datasets (56).

To further probe the relationship of ER, pS118-ER, and GRHL2 occupancy sites, we examined the signal intensity in the five binding categories. Binding signal of pS118-ER is substantially higher than ER sites, emphasizing the importance of this PTM in ER DNA binding (Fig. 2D, column 1, orange vs blue lines). However, the binding signal of pS118-ER and ER at regions co-occupied by GRHL2 did not appear noticeably different from regions where GRHL2 binding is absent (Fig. 2D, columns 1 and 2, solid vs dotted lines). Similarly, the presence of pS118-ER or ER does not substantially affect the binding signal of GRHL2 (Fig. 2D, column 3, solid vs dotted lines). These data indicate ER/pS118-ER binding is not preferentially stronger at GRHL2 binding sites and vice versa.

Previous studies showed GRHL2 can co-occupy chromatin with two factors that enhance ER transcriptional activity: the pioneer factor FOXA1 (58, 63, 91) and p300, a histone acetyltransferase that can deposit the enhancer histone mark H3K27ac (92, 93). We first asked whether the FOXA1 motif was enriched at pS118-ER/GRHL2 co-occupancy sites. While the FOXA1 motif has comparable enrichment across all categories with ER or pS118-ER binding, it is less prevalent at GRHL2-only binding sites (Fig. 3A), suggesting that FOXA1 could be preferentially recruited to sites occupied by both ER and GRHL2 over GRHL2 alone. Interestingly, the GRHL motif is the most prevalent motif at pS118-ER/GRHL2 and ER/GRHL2 co-occupancy sites while the ERE is most prevalent at pS118-ER-only and ER-only sites (Fig. 3A). Next, we used previously published FOXA1 (82) and p300 (83) ChIP-seq studies performed in E2-treated MCF7 cells (40 min and 1 hour treatments, respectively) to examine the presence of these two factors at pS118-ER/GRHL2 co-occupancy sites (Fig. 3B). For both factors, the binding signal was strongest at pS118-ER/GRHL2 co-occupancy sites followed by ER/GRHL2 co-occupancy sites and pS118-ER-only sites (Fig. 3B). Analysis of the active enhancer marks H3K27ac and H3K4me1 ((84), 30 min treatment), the promoter mark H3K4me3 ((84), 30 min treatment), and the repressive mark H3K27me3 ((85), 1 hour treatment) further revealed an enrichment of H3K27ac and H3K4me1 at pS118-ER/GRHL2 co-occupancy sites and ER/GRHL2 co-occupancy sites, relative to sites independently occupied by pS118-ER, ER, or GRHL2 alone (Fig. 3B). The promotor mark H3K4me3 is enriched at GRHL2 only sites relative to co-occupancy sites with pS118-ER and ER (Fig. 3B), which is consistent with the preferential localization of GRHL2 only sites in promotor regions compared to any other binding category categories (9.5% vs 2.5-5.5%) (Fig. 3B,C). The repressive H3K27me3 mark is weakest at pS118-ER/GRHL2 co-occupancy sites followed by ER/GRHL2 co-occupancy sites (Fig. 3B). These data indicate pS118-ER/GRHL2 co-occupancy sites are enriched at active enhancer regions.

**Figure 3:**
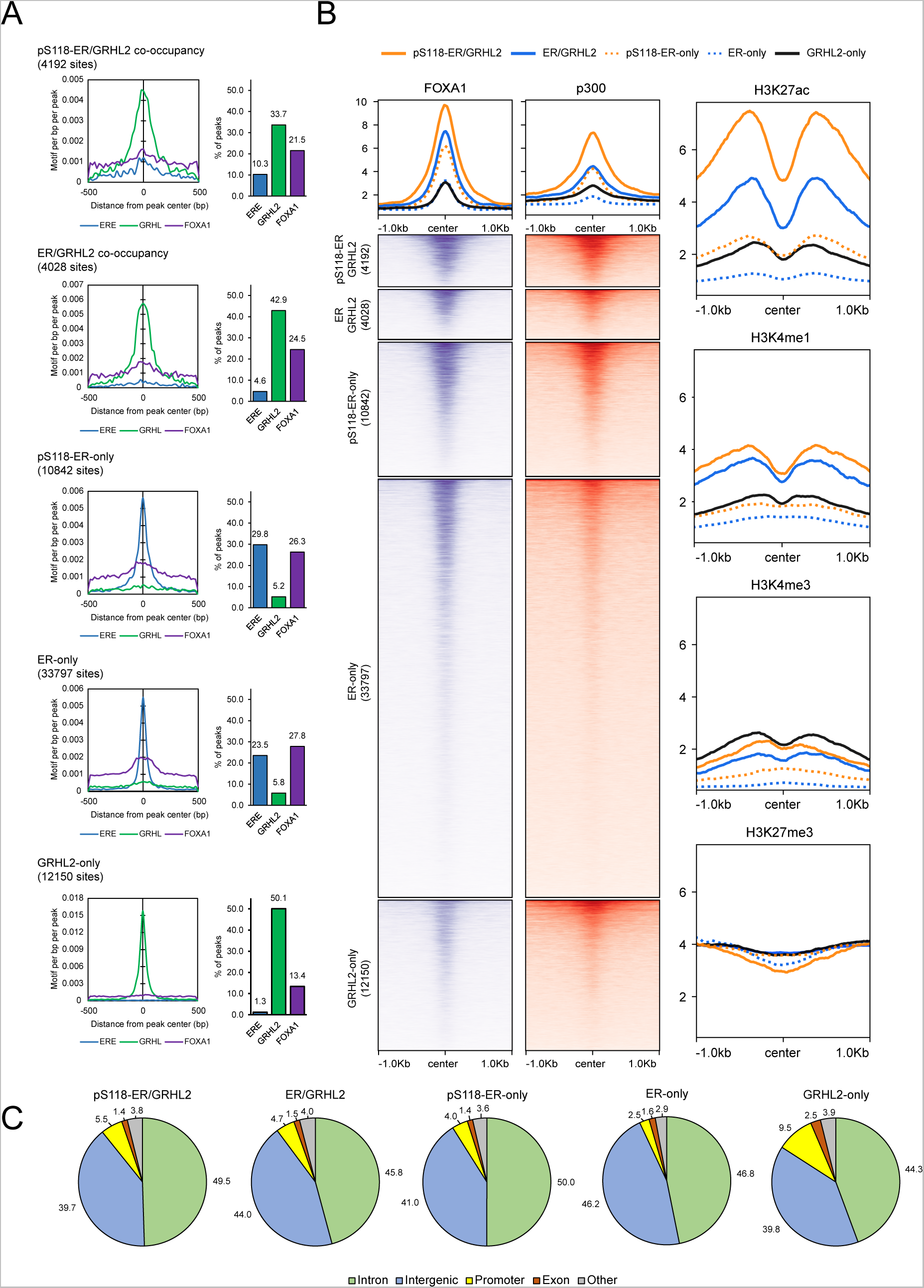
pS118-ER/GRHL2 co-occupancy sites are located at enhancers. A) Motif analysis of the GRHL, ERE, and FOXA1 motifs at co-occupancy sites. The first column shows enrichment of the indicated motif (ERE in blue, GRHL in green, FOXA1 in purple) relative to the peak center. The second column shows the percent of sites with each of the motifs. C) Heatmaps showing binding signal of FOXA1 and p300 and metaplots showing the signal of the histone modifications H3K4me1, H3K4me3, H3K27ac, and H3K27me3 at each of the previously established binding categories. Data were obtained from previously published ChIP-seq datasets (82–85). C) Genome location annotation of the five binding categories. Sites were annotated to introns, intergenic regions, promoters, exons, or other. Promoters were defined as -1kb to +100bp from the transcription start site.

### The loss of GRHL2 results in weaker pS118-ER DNA-binding

Given the preference of pS118-ER/GRHL2 co-occupancy sites for enhancer regions compared to either factor alone, we asked if GRHL2 is required for pS118-ER to bind to chromatin, and vice versa. First, we assessed the requirement of GRHL2 for pS118-ER chromatin occupancy by performing CUT&RUN for ER and pS118-ER in MCF7 cells depleted of GRHL2 using siRNA in the presence and absence of E2 (Fig. 4A). There was no change in pS118-ER induction by E2 after depletion of GRHL2 (Fig. 4A). High-confidence pS118-ER binding sites were identified by removing peaks that were detected in vehicle-treated cells and peaks that were not detected by the total ER antibody. In mock-transfected E2-treated cells (siCtrl), we identified 7,269 total ER and 2,860 pS118-ER binding sites. After the depletion of GRHL2, 4,171 ER sites were lost (57%) and 1,089 ER sites were gained. 3,098 ER sites were maintained in both control and GRHL2-depleted conditions (Fig. 4B,C). Both total ER and pS118-ER binding signals were substantially decreased in GRHL2-depleted cells, including instances when ER binding was maintained in both control and GRHL2-depleted conditions (Fig. 4C). Phosphorylation of S118 could be detected in control conditions at 1,178 (28.2%) of the ER binding sites that were lost and 2,026 (65.4%) of ER binding sites that were maintained in both control and GRHL2-depleted conditions (Fig. 4B,C). However, only 127 (11.7%) of new ER binding sites identified in GRHL2-depleted conditions were phosphorylated at S118 (Fig. 4B,C). These data indicate GRHL2 is required for robust pS118-ER DNA occupancy.

**Figure 4:**
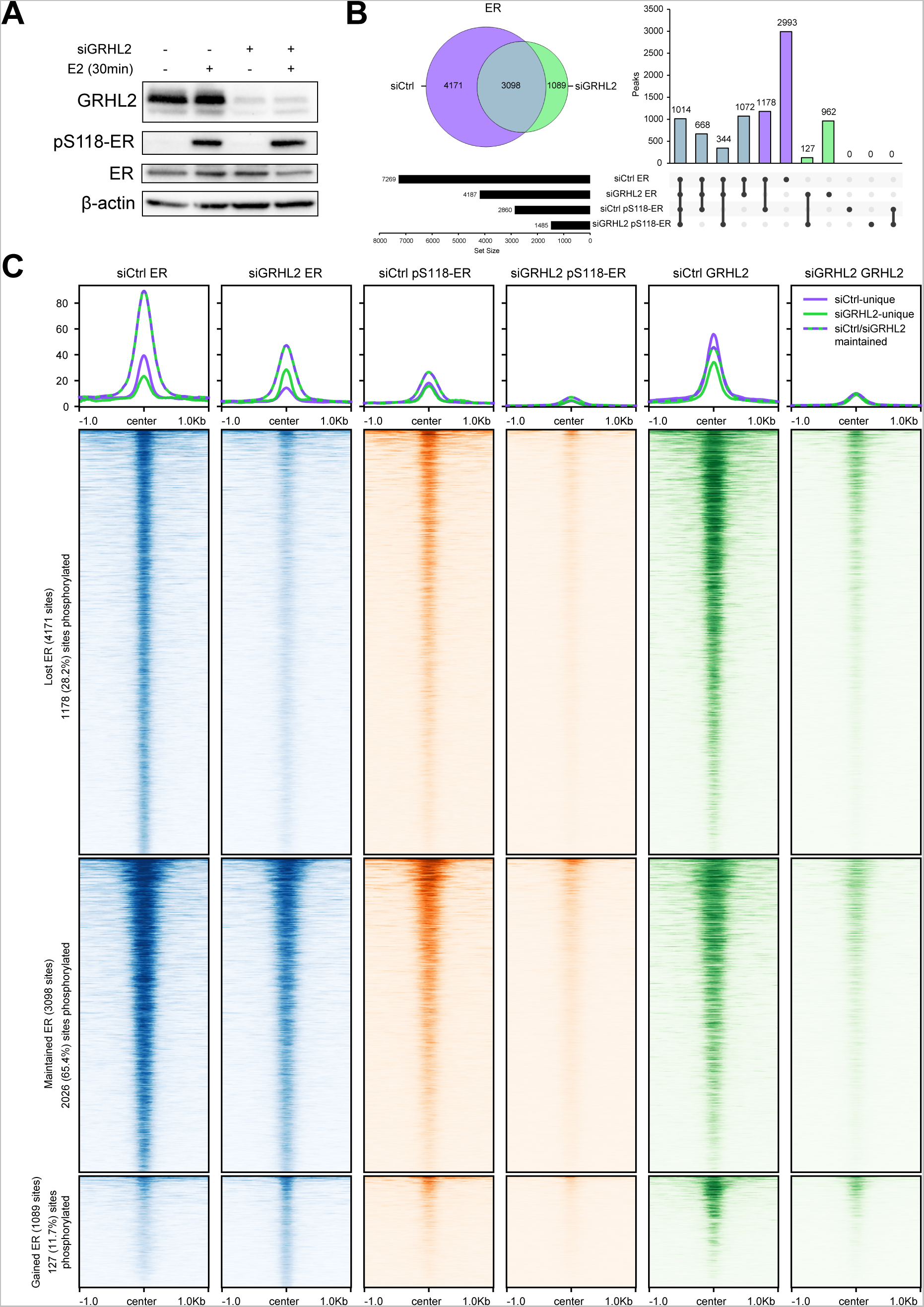
The loss of GRHL2 causes a reduction in ER and pS118-ER chromatin binding. A) Representative western blot of MCF7 samples used in CUT&RUN experiments. Cells were transfected with siCtrl or siGRHL2 and treated for 30 minutes with vehicle or 10 nM E2. β-actin is used as a loading control. B) Venn diagram and upset plot indicating the overlap of ER and pS118-ER CUT&RUN peaks in E2-treated, siCtrl- or siGRHL2-transfected cells. C) Heatmaps showing CUT&RUN signal for ER, pS118-ER, and GRHL2 at lost ER sites (purple), maintained ER siCtrl/siGRHL2 sites (purple and green dashed), and gained ER sites (green). The percent of ER sites detected by the pS118-ER antibody are indicated in region labels.

Conversely, to determine if the phosphorylation of S118 on ER is required for GRHL2 DNA-binding, we performed ChIP-seq for GRHL2 in two independent MCF7 CRISPR knock-in cell lines in which the serine at position 118 has been mutated to an alanine (S118A-ER), thereby preventing phosphorylation at this site (Fig. 5A). In wild-type (WT) cells with intact pS118-ER, GRHL2 occupied 16,123 sites. In knock-in cell lines with S118A-ER, GRHL2 occupied 14,682 sites in one clone (S118A #1), and 16,634 sites in the second clone (S118A #2). The majority of binding sites (13,066) were shared among the three cell lines (Fig. 5B). Further, GRHL2 DNA-binding signal in the two S118A mutants was comparable to the WT cells. Together, these data indicate that pS118-ER is not reciprocally required for GRHL2 to bind DNA (Fig. 5C).

**Figure 5:**
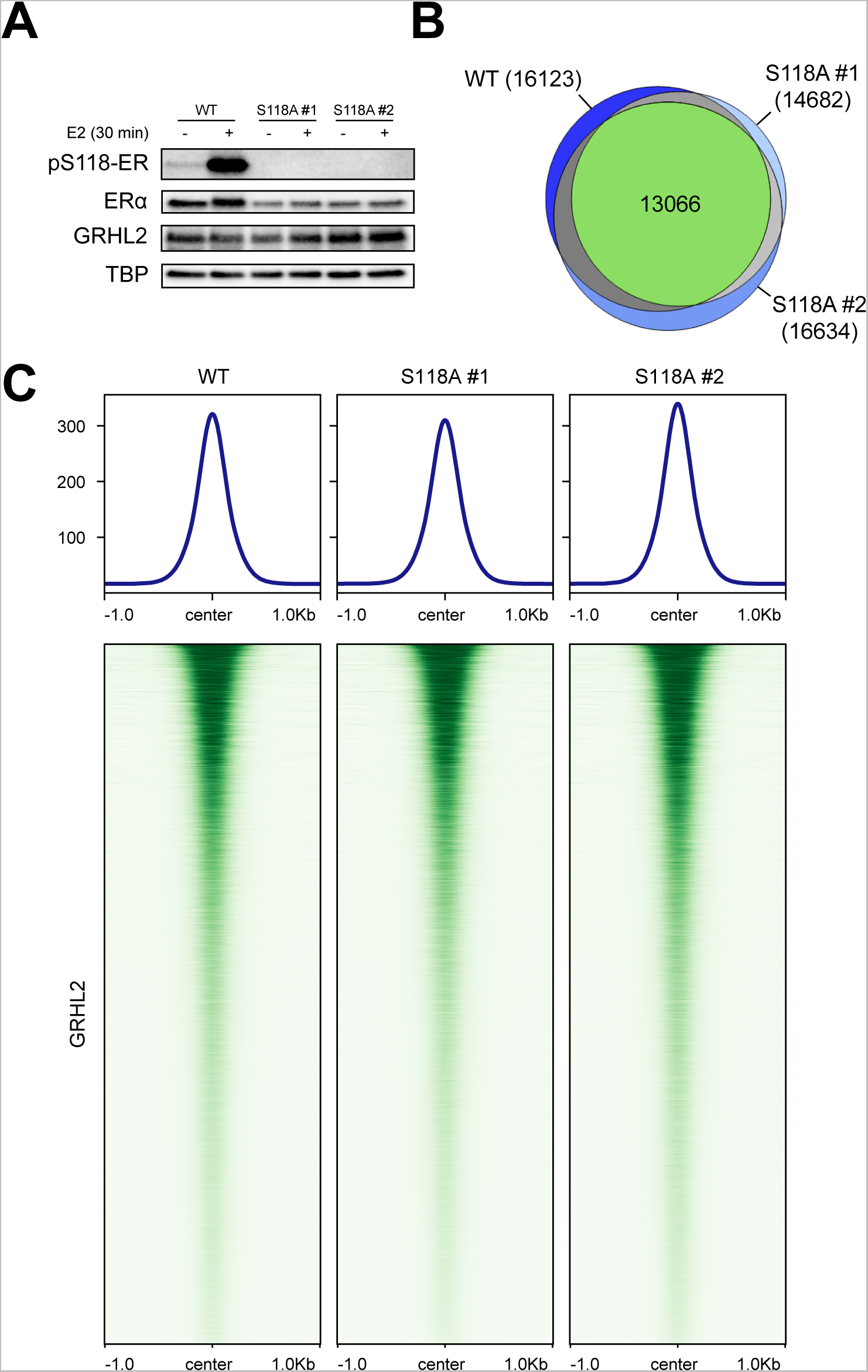
Loss of pS118-ER does not affect GRHL2 chromatin binding. A) Representative western blot of WT and S118A mutant MCF7 cells used for ChIP-seq. Cells were treated for 30min with vehicle or 10 nM E2. TBP is used as a loading control. B) Venn diagrams showing the overlap of E2 GRHL2 binding sites between WT cells and two S118A mutant clones. C) Heatmaps showing GRHL2 binding signal at collective GRHL2 binding sites across all three cell lines.

### GRHL2 and estrogen co-regulated genes are involved in cell migration

After establishing that pS118-ER/GRHL2 co-occupancy sites bind preferentially in enhancer regions and that GRHL2 is necessary for pS118-ER chromatin binding, we next investigated what genes and biological processes these two factors co-regulate. To identify these genes, we performed RNA-seq in triplicate in E2-treated MCF7 and T47D cells which had been depleted of GRHL2 using siRNA (Fig. 6A, Supplemental Fig. 1A). E2 activation was confirmed by the induction of pS118-ER (12, 19) and downregulation of total ER levels (94) (Fig. 6A, Supplemental Fig. 1A). As before, the loss of GRHL2 did not affect E2-mediated pS118-ER induction (Fig. 6A, Supplemental Fig. 1A). Principle component analysis (PCA) showed a large amount of variance between vehicle and E2 treatments in both cell lines, with a larger variance observed in MCF7 cells (Fig. 6B, Supplemental Fig. 1B). There was a larger variance between siCtrl and siGRHL2 in T47Ds than in MCF7s (Fig. 6B, Supplemental Fig. 1B). Differential expression analysis was next performed using DESeq2 and the results were filtered for genes with an adjusted p-value of less than 0.05 and a log_2_ fold change of greater than 1 or less than -1. Genes were defined as primarily controlled by E2-activated ER (ER-only), primarily controlled by GRHL2 (GRHL2-only), or co-regulated by both ER and GRHL2 (ER/GRHL2 co-regulated) (Fig. 6C, Supplemental Fig. 1C, Supplemental Tables 1 and 2).

**Figure 6:**
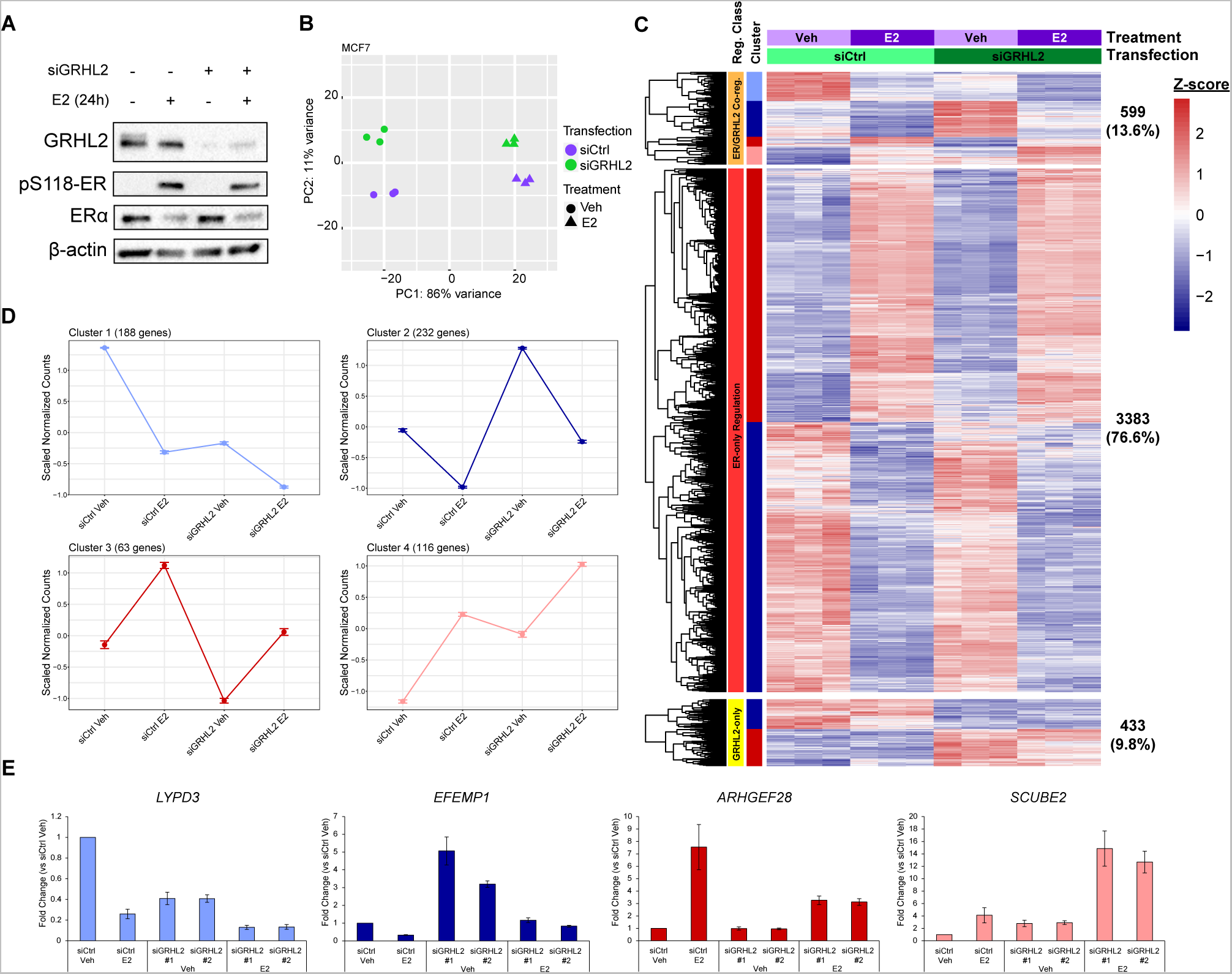
GRHL2 can enhance and antagonize E2-mediated ER transcriptional activity. A) Representative Western blot of MCF7 samples used in RNA-seq. Cells were transfected with siCtrl or siGRHL2 and treated for 24 hours with vehicle or 10 nM E2. β-actin is used as a loading control. B) PCA plot of MCF7 RNA-seq data displaying differences between siCtrl (purple), siGRHL2 (green), vehicle treatment (circles), and E2 treatment (triangles). C) Heatmap showing the expression of ER/GRHL2 co-regulated genes, ER-only regulated genes, and GRHL2-only regulated genes. D) Gene expression profiles of ER/GRHL2 co-regulated gene clusters. Error bars represent mean ± standard error. E) qPCR validation of selected genes from each of the ER/GRHL2 co-regulated gene clusters using two different siRNAs against *GRHL2*. All values are relative to siCtrl Veh. Error bars represent mean ± standard error.

Examination of the ER/GRHL2 co-regulated genes revealed dual roles of GRHL2 in the regulation of E2-mediated ER transcriptional activity. ER/GRHL2 co-regulated genes can be broken down into four clusters (Fig. 6C,D, Supplemental Fig. 1C,D). When GRHL2 is lost, genes in clusters 1 and 4 show increased downregulation and upregulation by estrogen, respectively. These data indicated that when GRHL2 is present it opposes the regulatory action of ER on these genes. Genes in cluster 2 and 3 show decreased downregulation and upregulation by ER when GRHL2 is lost. At these genes, when GRHL2 is present it enhances ER transcriptional regulation. Using qRT-PCR and two independent siRNAs against *GRHL2,* we validated these co-regulation patterns in a selection of genes: *LYPD3* (cluster 1), *EFEMP1* (cluster 2), *ARHGEF28* (cluster 3), and *SCUBE2* (cluster 4) (Fig. 6E). Notably, *LYPD3* was previously identified as a GRHL2 target gene in a tamoxifen resistant breast cancer cell line (63). These data demonstrate that GRHL2 has pleiotropic effects on E2-induced pS118-ER transcriptional activity and can work as either a transcriptional activator or repressor to influence the magnitude of estrogenic responses in a gene-specific manner.

To determine which biological processes that ER/GRHL2 co-regulated genes may influence, we performed gene ontology analysis using clusterProfiler. As expected, E2-only genes were primarily associated with mitosis and cell cycle-related pathways (95) and GRHL2-only genes annotated to epithelial terms such as “regulation of cell-cell adhesion” (24, 27, 31) and “wound healing” (96) (Supplemental Table 3). Gene ontology analysis of ER/GRHL2 co-regulated genes found four of the top ten most significant pathways in MCF7s were related to cell migration and motility (Fig. 7A, Supplemental Table 3). To test the effect of E2-activate ER and GRHL2 on migration, we took advantage of MCF7 cell pools which have undergone CRISPR-mediated depletion of GRHL2 (GRHL2-KO) (Fig. 7B-D). Cells were treated with vehicle or E2 for 24 hours prior to the creation of a cell-free gap. GRHL2-KO cells migrated across the cell-free gap more quickly than wild type (WT) cells over the course of 24 hours, and migration was further enhanced by the addition of E2. We determined the increase in migration was not a result of increased growth based on cell cycle analysis which indicated the loss of GRHL2 did not impact E2-induced growth responses in this time frame (Fig. 7E). These data indicate a co-regulatory role for GRHL2 and E2-induced pS118-ER in regulating cell migration. In addition, these data demonstrate that GRHL2 is a critical blockade that inhibits the migratory properties of MCF7 cells in the presence of activated ER.

**Figure 7:**
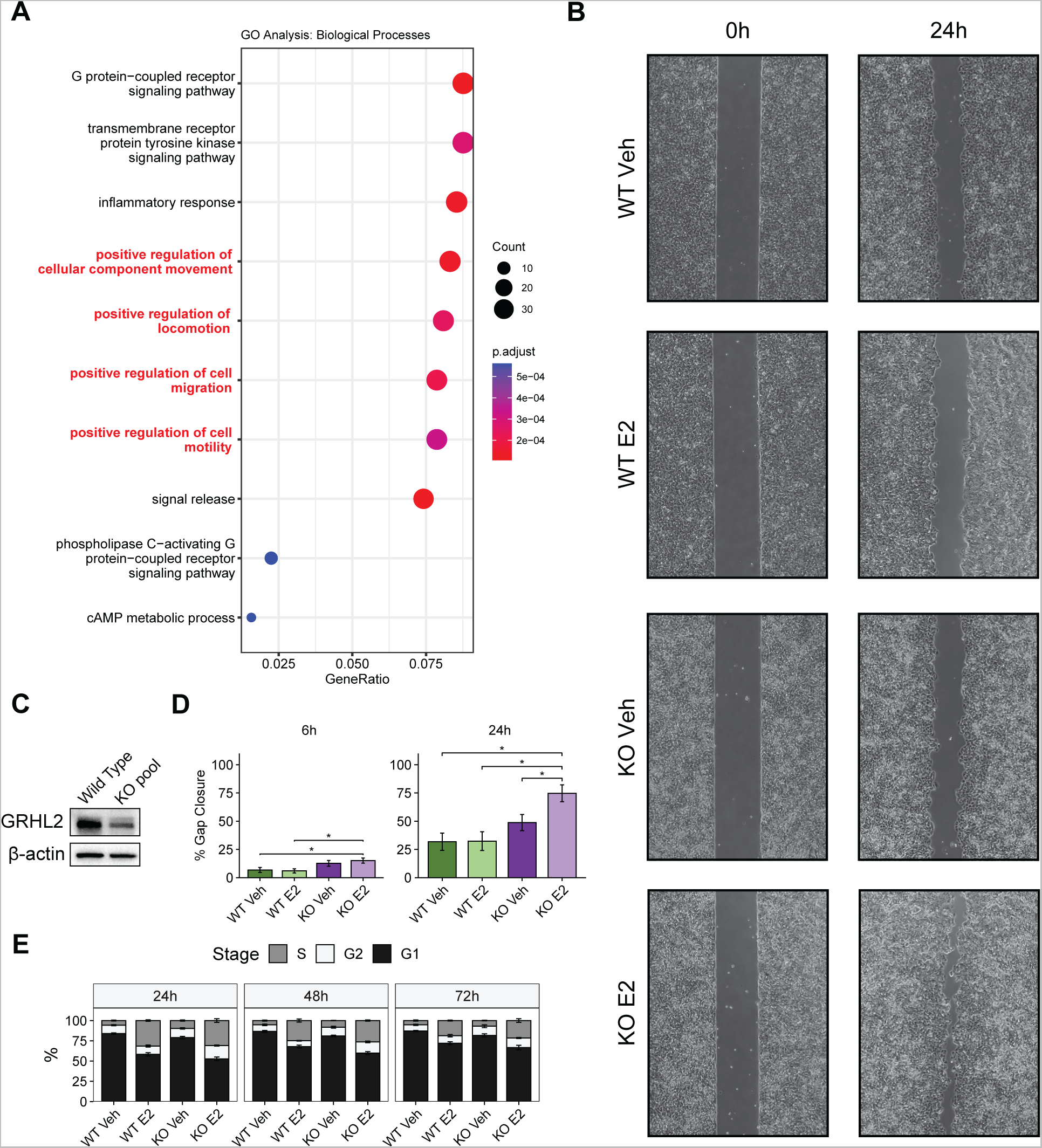
GRHL2 and E2 co-regulate migration. A) Gene ontology analysis for biological processes performed with clusterProfiler on ER/GRHL2 co-regulated genes. Terms related to cell migration, motility, and locomotion are highlighted in red. The size of the dots represents the number of genes annotated to a given term, and color represents the adjusted p-value with red being the most significant. B) Brightfield images of WT and GRHL2-KO cell migration across a cell-free gap over the course of 24 hours. C) Representative Western blot showing the loss of GRHL2 in pooled CRISPR GRHL2-KO cells. D) Quantification of the percent gap closure over 6 hours and 24 hours. Error bars represent mean ± standard error and * indicates a p-value < 0.05. E) Flow cytometry analysis showing the percent of cells in each of the cell cycle stages (S, G2, G1). WT and GRHL2-KO cells were treated with vehicle or 10 nM E2 over 24h, 48h, and 72h. Error bars represent mean ± standard error.

## DISCUSSION

Phosphorylation of S118 on the estrogen receptor is necessary for maximal ER-transcriptional activity and chromatin-binding (12, 17, 19, 97). Our group identified a specific enrichment of the GRHL2 motif near pS118-ER occupancy sites suggesting an important role for GRHL2 in pS118-ER activity (19). In this report we examined the co-dependency of GRHL2 and pS118-ER chromatin-occupancy and transcriptional regulation of gene expression. We found that the loss of GRHL2 results in a global decrease in pS118-ER chromatin-occupancy, demonstrating a critical role for GRHL2 in promoting maximal pS118-ER recruitment to chromatin. GRHL2 can bind to chromatin prior to ligand treatment and pS118-ER/GRHL2 co-occupancy sites tend to be localized to active enhancer regions marked by H3K27ac and H3K4me1. Transcriptionally, GRHL2 can both enhance and antagonize E2-induced pS118-ER transcriptional activity, indicating a role for GRHL2 in tuning pS118-ER-mediated gene expression. These studies lead us to propose a model in which GRHL2 primes chromatin prior to ligand treatment to facilitate the recruitment of pS118-ER and then further fine tunes E2-mediated pS118-ER transcriptional activity through the recruitment or repression of additional coregulators (Fig. 8). The full range of coregulators GRHL2 recruits or interacts with are still unknown and may vary based on the genetic background of a given cell.

**Figure 8:**
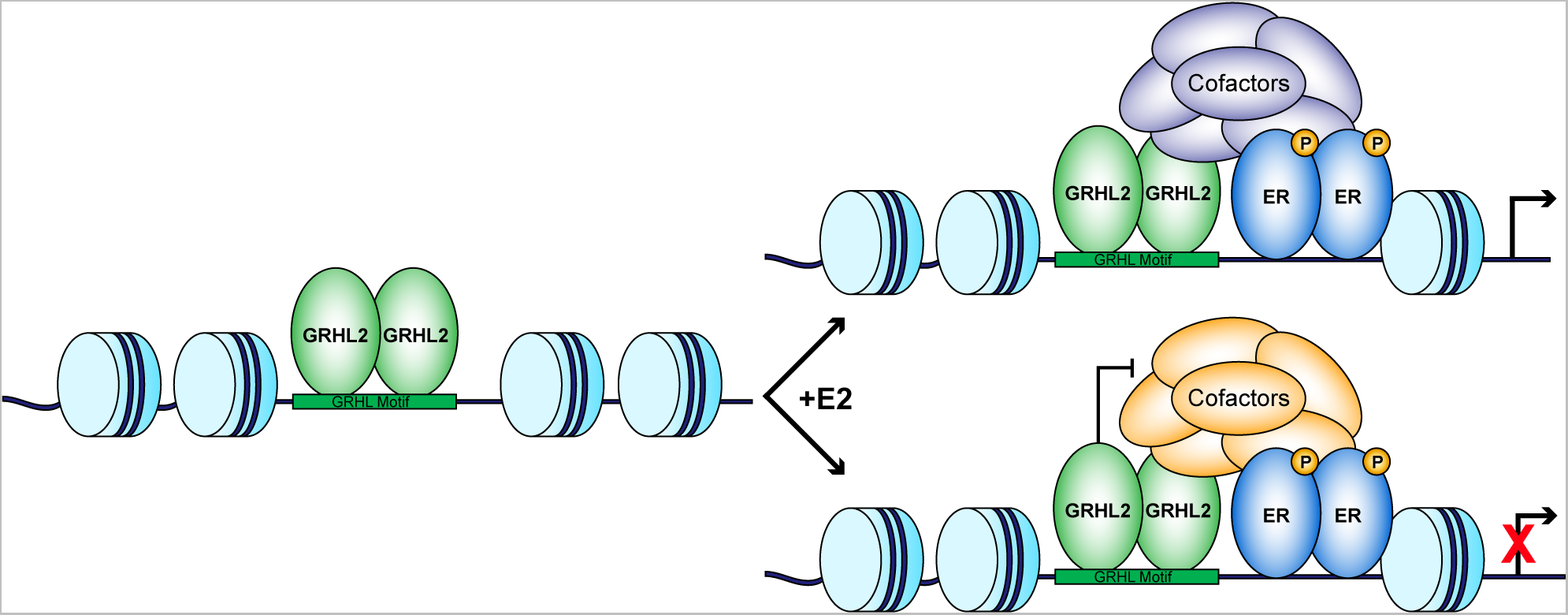
Model of GRHL2 function in facilitating pS118-ER binding and transcriptional activity. GRHL2 binds to the GRHL motif in enhancers prior to ligand treatment. Upon exposure to E2, GRHL2 can either promote gene transcription by facilitating the recruitment of coregulators like MLL3, or repress gene transcription by repressing coregulators, like p300.

GRHL2 occupies sites destined to be co-bound by ER prior to E2-mediated ER activation, and GRHL2 is necessary for robust pS118-ER DNA binding. In contrast, a S118A mutation in ER that prevents phosphorylation failed to alter GRHL2 binding. These data suggest that GRHL2 primes enhancers for ER binding events. Similar observations were made in studies of the highly conserved *Drosophila melanogaster* homolog grainyhead (Grh) (98–101). These data have led to the hypothesis that the GRHL family may function as pioneer factors, a specialized class of transcription factors that influence transcriptional activity by promoting an open chromatin structure which promotes the recruitment of other transcription factors (102, 103). Decreased Grh in the developing *D. melanogaster* and decreased GRHL2 in epiblast-like cells resulted in a reduction of global chromatin accessibility (98, 99, 104). The same was observed when all three mammalian GRHL family members were knocked down in MCF7 cells (98). Another study using transgenic GFP reporters found Grh was bound to and promoted the accessibility of various enhancer regions prior to their activation (98). Further, mutation of the Grh motif at a selection of enhancers resulted in reduced chromatin accessibility (98). Interestingly, while the ERE is typically the most highly enriched motif at pS118-ER occupancy sites (19), we instead discovered a high prevalence of the GRHL motif at pS118-ER/GRHL2 co-occupancy sites. These data together suggest that the presence of the GRHL motif may identify a select set of ER enhancers that require chromatin opening in order for pS118-ER to bind DNA.

Previous studies have found GRHL2 can specifically associate with open and active enhancer regions marked by H3K27ac and H3K4me1 (56, 104). In agreement, we observed an association of GRHL2 binding sites with H3K27ac and H3K4me1 in breast cancer cells, particularly when it co-occupied chromatin with ER or pS118-ER. A previous study examining co-occupancy sites of GRHL2 and androgen receptor (AR), another nuclear receptor, also found an enrichment of H3K27ac at co-occupancy sites compared to either factor alone (35). These data suggest that other transcription factors and coregulators are needed in addition to GRHL2 to fully engage active enhancers. We also found the strongest enrichment of FOXA1 binding signal at pS118-ER/GRHL2 co-occupancy sites. FOXA1 frequently binds in active enhancer regions marked by H3K4me1 (62) and is known to facilitate ER recruitment to chromatin (105). GRHL2 has been shown to work in concert with FOXA1 to recruit the histone methyltransferase MLL3 which in turn deposits H3K4me1 (58). GRHL2 can also directly interact with MLL3 via two amino acids in the GRHL2 transactivation domain, suggesting it may directly recruit MLL3 (106). However, regardless if MLL3 is present, GRHL2/FOXA1 co-occupancy sites also show stronger enrichment of H3K4me1 than sites where either factor occupies chromatin individually (58). This suggests that other coregulators may also contribute to H3K4me1 deposition at these sites. Overall, our data provide additional evidence that the combination of GRHL2, FOXA1, and pS118-ER more faithfully mark active enhancers than the individual factors.

Given the association of pS118-ER with upregulated genes (19) and the preference of GRHL2 and pS118-ER for active enhancers, we were surprised to find in our transcriptomic analysis that GRHL2 can both enhance and antagonize E2-mediated gene expression. This suggests that GRHL2 may interact with other coregulators to modulate pS118-ER transcriptional function. As discussed above, these factors likely include FOXA1 and MLL3, which are associated with marks of active enhancers. In this report we found pS118-ER/GRHL2 co-occupancy sites display the strongest signal for the ER co-activator p300 (93, 107), a histone acetyltransferase which can deposit the active enhancer mark H3K27ac (108). Although p300 could be responsible for the presence of H3K27ac at pS118-ER/GRHL2 co-occupancy sites, GRHL2 can inhibit p300 activity via an interaction of a 13 amino acid segment of the GRHL2 DNA binding domain with the C-terminal end of p300 (92). Based on this finding, it is possible that GRHL2 could inhibit p300/ER transcriptional complexes leading to repression at select genes. Therefore, one explanation of how GRHL2 can either activate or repress pS118-ER gene transcription is via GRHL2 activity on other transcription factors (FOXA1) or histone modifiers (MLL3, p300) that make up the pS118-ER transcriptional complex.

In addition to exploring the interplay between GRHL2 and pS118-ER in chromatin binding, we also wanted to understand what biological processes are co-regulated by GRHL2 and E2-activated pS118-ER. As expected, genes associated with cell cycle and epithelial cell integrity aligned with ER-only and GRHL-only gene sets respectively, consistent with their well-described roles (24, 27, 31, 95, 96). Gene ontology analysis of ER/GRHL2 co-regulated genes, however, showed a large proportion of co-regulated genes were involved in migration. We subsequently found that the loss of GRHL2 can increase cell migration and this effect is further enhanced by the addition of E2. A previous study in ER-negative cell lines also found GRHL2 influences migration in a tumor suppressive manner (40), but this study is the first to show the co-regulatory role of GRHL2 and ER in this phenotype. In the hormone-dependent ER-positive breast cancer model tested here, GRHL2 serves as an important blockade that inhibits cell migration. An example of an ER/GRHL2 co-regulated gene that was annotated to cell migration in the gene ontology analysis was *SASH1*, a tumor suppressor that inhibits invasion and migration (109, 110). *SASH1* regulation is described by cluster 1, in which E2 downregulates *SASH1* and GRHL2 depletion results in further downregulation of *SASH1* expression. Regulation of *SASH1* may be an example of an instance where GRHL2 depletion results in an expression threshold being crossed and factors like SASH1 can no longer enact their tumor suppressive action, resulting in increased cell migration.

In summary, we have established a role for GRHL2 in facilitating the binding of pS118-ER to DNA and have shown that sites co-occupied by GRHL2 and pS118-ER identify a set of enhancers marked by H3K27ac and H3K4me1. We propose a model for the potential interactions among GRHL2, pS118-ER, and other coregulators like p300 and FOXA1 to either complement or oppose pS118-ER transcriptional activity. Finally, our data indicate that decreases in GRHL2 lead to increased cell migration that is further enhanced by estrogen stimulation. These data identify dual activities of GRHL2 and its modulation of activated ER and reveal a complexity to the interplay between GRHL2 and pS118-ER that impacts cancer phenotypes, particularly migration.

## Supporting information

Supplemental Table 1 - MCF7 RNA-seq data

Supplemental Table 2 - T47D RNA-seq data

Supplemental Table 3 - MCF7 GO BP

## ACKNOWLEDGEMENTS

We thank Kathryn Fox and the University of Wisconsin Carbone Comprehensive Cancer Center Flow Cytometry Laboratory for their assistance with proliferation studies. We also thank Dr. David Lung for his input into the manuscript. This work is supported by NIH grant R01 CA260140 to ETA.

## FIGURE LEGENDS

**Supplemental Figure 1: Similar patterns of GRHL2 regulation of E2-mediated ER transcriptional activity are observed in T47D cells.** A) Representative Western blot of T47D samples used in RNA-seq. Cells were transfected with siCtrl or siGRHL2 and treated for 24 hours with vehicle or 10 nM E2. β-actin is used as a loading control. B) PCA plot of T47D RNA-seq data displaying differences between siCtrl (purple), siGRHL2 (green), vehicle treatment (circles), and E2 treatment (triangles). C) Heatmap showing the expression of ER/GRHL2 co-regulated genes, E2-only regulated genes, and GRHL2-only regulated genes. D) Gene expression profiles of ER/GRHL2 co-regulated gene clusters. Error bars represent mean ± standard error.

