## Supplemental Figure 1 for "GRHL2 enhances phosphorylated estrogen receptor DNA-binding and regulates ER-mediated transcriptional activation and repression"

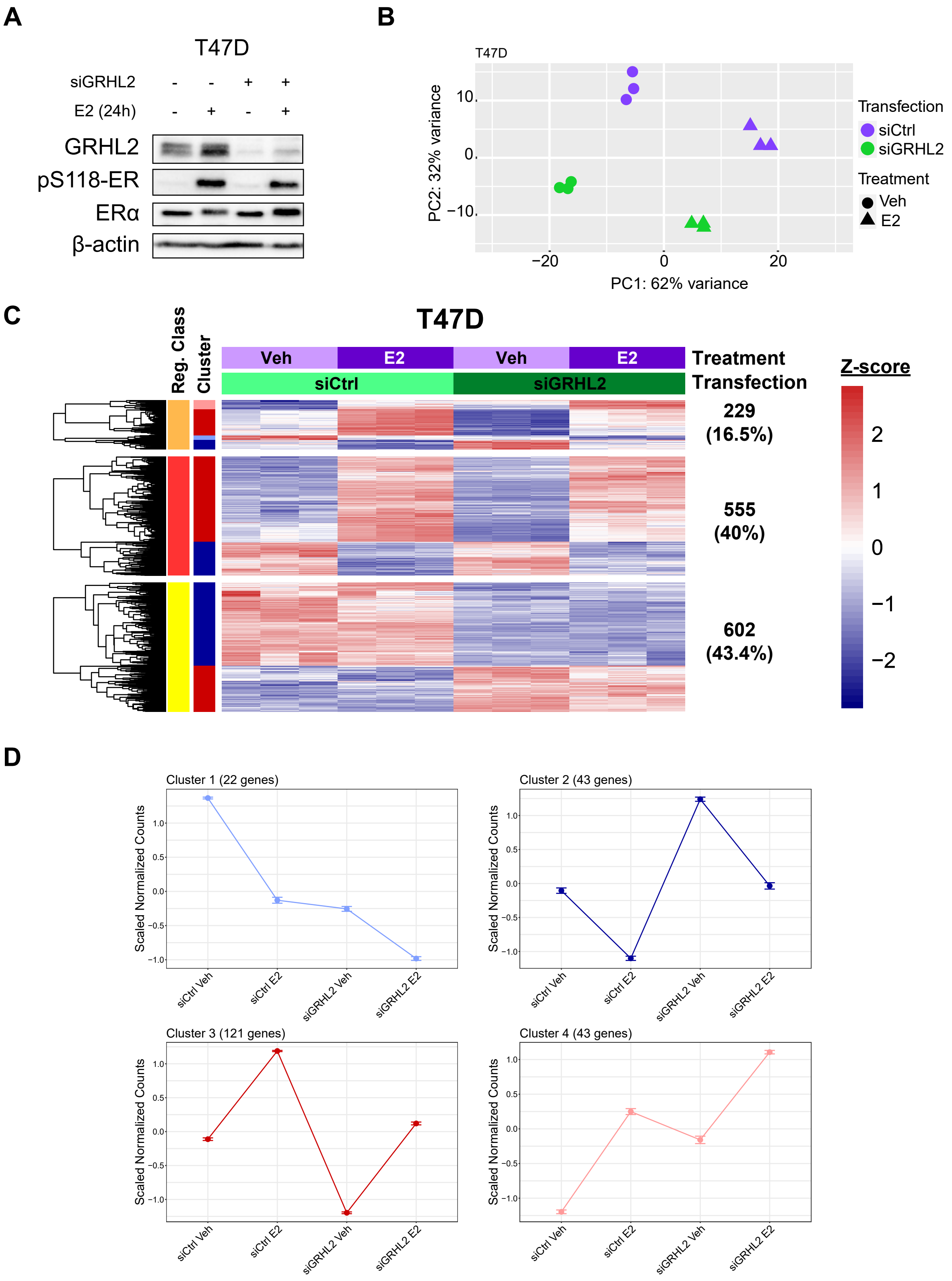


**Supplemental Figure 1: Similar patterns of GRHL2 regulation of E2-mediated ER transcriptional activity are observed in T47D cells.** A) Representative Western blot of T47D samples used in RNA-seq. Cells were transfected with siCtrl or siGRHL2 and treated for 24 hours with vehicle or 10 nM E2. β-actin is used as a loading control. B) PCA plot of T47D RNA-seq data displaying differences between siCtrl (purple), siGRHL2 (green), vehicle treatment (circles), and E2 treatment (triangles). C) Heatmap showing the expression of ER/GRHL2 co-regulated genes, E2-only regulated genes, and GRHL2-only regulated genes. D) Gene expression profiles of ER/GRHL2 co-regulated gene clusters. Error bars represent mean ± standard error.
